# Generalizing Drift Diffusion Models with a State-Dependent Framework

**DOI:** 10.1101/2025.11.05.686838

**Authors:** Juan Sebastian Cely-Acosta, Pete C Trimmer

## Abstract

Decision-making in natural environments often involves a trade-off between speed and accuracy, particularly in high-stakes scenarios like predator-prey interactions. Drift Diffusion Models (DDMs) have been used extensively to study speed-accuracy trade-offs, but such work has typically assumed that decisions are made independently of one another. In the wild, the consequences of one decision will often influence future choices. To address this gap, we introduce a new framework – State-Dependent Drift Diffusion Models (S3DMs) – which integrate state-dependent variables into the decision-making process. Using simulations of foraging scenarios, we demonstrate that S3DMs can predict markedly different behavioural outcomes compared to a standard DDM. By incorporating sequential and interdependent decisions, S3DMs offer a framework that is arguably both more realistic and more biologically relevant. Our research contributes to integrating functional and mechanistic explanations of behaviour and suggests a promising direction for future studies in behavioural ecology and neuroscience.

## Introduction

Survival often depends on the ability to make rapid decisions; for instance, whether a stimulus should be responded to as though it is a predator or prey. Signal Detection Theory (SDT; Green & Swets, 1966; Haselton & Nettle, 2006) and state-dependent variants (Ehlman et al., 2019; Trimmer et al., 2017) have been used to represent such scenarios. However, signal detection models typically assume that decisions are made instantaneously and thus overlook the mechanisms that control decision time (Pleskac & Busemeyer, 2010; Wixted, 2020). The sensory evidence received by an organism is frequently noisy, which means that signals are not always reliable, and it could be better to wait for more evidence to execute an appropriate action. How quickly a decision is made will typically govern the consequences of an action, so it is necessary for an organism to wait only what is optimal to guarantee its survival.

Sequential-sampling models were developed based on the understanding that an organism’s response to a repeated stimulus can vary, even under seemingly identical conditions, both in the choice of action and in the time taken to respond (Heitz, 2014; Smith & Ratcliff, 2024). A major subclass of these models, known as random walk models, is widely used in psychology to analyse decision-making and response times (Smith & Ratcliff, 2015). These models are grounded in the Sequential Probability Ratio Test (SPRT), developed by Wald (Bogacz et al., 2006). SPRT operates as a dynamic process, continuously evaluating the likelihood of two competing hypotheses as new data becomes available (Gold & Shadlen, 2007; Smith & Ratcliff, 2015). The likelihood ratio is updated with each piece of incoming data, and a decision is made in favour of one hypothesis once the ratio surpasses a pre-defined threshold (Smith & Ratcliff, 2015). Within this framework, decision-making is viewed as a statistical process where noisy evidence or stimulus information accumulates until a response criterion (i.e., threshold) is reached (Smith & Ratcliff, 2015). The evidence is considered noisy due to either variability in the stimulus or noise at the neural level (Smith & Ratcliff, 2024). Importantly, these models assume that evidence is accumulated in discrete time-steps, and in this process, evidence for one response simultaneously acts as evidence against the other, represented by a single fluctuating sum that changes as evidence is sampled (Ratcliff et al., 2016).

If we take the SPRT and allow the time between updates to decrease to zero and maintain that the evidence received per unit of time (i.e., drift rate) is constant and normally distributed, then the resulting process is a Brownian motion with drift, also known as a Wiener process (Smith & Ratcliff, 2024), or a drift diffusion model (DDM) (Bogacz et al., 2006). The decision-making process in a DDM is similar to the SPRT model, where evidence for one response option drives the process in an upward direction, closer to the upper threshold, while evidence for the other response option drives it in a downward direction (Trimmer et al., 2013). This process continues until one of the thresholds is reached, at which point the corresponding decision is reached. The crucial difference is that in a DDM evidence accumulates continuously rather than at discrete points in time (Smith & Ratcliff, 2024).

DDMs have been extensively used to describe behavioural data coming from a variety of laboratory tasks that require an individual to make a fast two-choice decision (Ratcliff et al., 2016). These models assert that successive samples of noisy evidence coming from a stimulus are accumulated in the organism favouring either one of the possible options until at some point enough evidence in favour of one option is accumulated and a response is made accordingly (Smith & Ratcliff, 2024). Importantly, diffusion models have been effectively linked to neurophysiological measures for single cells and populations of neurons in rodents, monkeys and humans, showing correspondence with the underlying biological mechanisms (Brunton et al., 2013; Roitman & Shadlen, 2002; Bode et al., 2012; Polanía et al., 2014; Ratcliff et al., 2003, 2007; Smith & Ratcliff, 2004; Ratcliff & McKoon, 2008; Van Vugt et al., 2012).

As with standard SDT (Egan, 1975), DDMs typically assume that each decision is independent of one another (Nguyen et al., 2019; Griffith et al., 2021). However, when it comes to organisms in their natural environments, the consequences of a decision can influence further decisions (Ehlman et al., 2019; Lepora & Pezzulo, 2015). Thus, taking these influences into account could enhance the realism of our models (Houston et al., 2024). Moreover, this interdependence among decisions might also result in counterintuitive effects, leading to predictions that diverge from those of standard models (Ehlman et al., 2019; McNamara & Trimmer, 2019).

In this paper, our aim is to develop a more realistic family of DDMs using a state-dependent modelling framework, which we call state-dependent drift diffusion models (S3DMs). We compare the behavioural predictions of the S3DMs with those of a standard DDM in natural foraging scenarios where environmental risks, such as predation, are present and responses need to vary according to the level of risk.

## Methods

We consider a simple foraging scenario and conduct simulations to compare the outcomes of a standard DDM and the new S3DMs. Our primary focus is on the models’ predictions regarding how variations in danger levels, specifically predator prevalence, influence the behavioural responses of an organism. In this section, we first describe the general foraging scenario and the application of a standard DDM to it. Next, we explain the rationale behind state-dependent modelling and its role in the development of the S3DMs.

### Drift diffusion model

We are interested in how environmental cues indicating threats (e.g., predators) or opportunities (e.g., food) influence decision-making in organisms that occupy intermediate trophic levels. We analyse a foraging scenario in which an individual needs to respond to a novel stimulus in the form of another animal that could be either predator or prey. For simplicity, we will disregard the presence of conspecifics or other stimuli that are neither predators nor prey. The focal individual needs to approach prey fast to catch it while remaining vigilant of predators to react quickly and avoid them. Thus, the individual faces a speed-accuracy trade-off (Chittka et al., 2009). In a DDM, the trade-off is controlled by the position of the decision thresholds; more distant thresholds result in slower but more accurate decisions (Bogacz, 2007).

We employ a DDM similar to one previously used in research on evolutionary approaches to mood (Trimmer et al., 2013). In this model, the payoffs for actions are linked to the probability of successfully capturing prey and the risk of being caught by a predator. We assume that the probability of the environmental condition (predator present or not) can be estimated; we denote the probability of predator being present as *p*, so (1 – *p*) is the probability that the encountered individual is prey. When an animal appears, the DDM determines how ready the individual should be to decide whether to approach or avoid the encountered animal.

We assume that the drift is positive when the novel stimulus comes from a prey and negative when it comes from a predator. The two thresholds represent an approaching response (upper threshold) and an avoidance response (lower threshold). These thresholds are influenced by the payoffs associated with each choice and how these payoffs change with time. When the payoffs for each situation are known, the optimal thresholds, which trade-off speed and accuracy to maximize expected reward, can be calculated. From these thresholds, it is possible to determine the expected decision times and accuracies.

The decision time, *T*, will typically depend on the decision that is reached. We denote the time taken to decide to approach by *T*_*ap*_, and the time taken to decide to avoid by *T*_*av*_.

The optimal thresholds depend on the payoffs, and these payoffs in turn depend on the reproductive value. Reproductive value, denoted by *V*, reflects the individual’s potential for future reproductive success (Trimmer et al., 2013; Fisher, 1930). Table 1 shows the payoff matrix with the payoff for each combination of response and circumstance. The payoffs depend on the circumstance, response, time to decide on that response, and reproductive value.

**Table 1.** Payoff matrix for the DDM.

|  |  | Response |  |
| --- | --- | --- | --- |
|  |  | Approach | Avoid |
| Environmental condition | Predator | 0 | $S(T_{av})V$ |
| | Prey | $C(T_{ap})(V + m) + (1 - C(T_{ap}))V$ | $V$ |

We assume that the probability of escaping from a predator, *S(T*_*av*_), decreases with time to reach the decision, *T*_*av*_, according to *S(T*_*av*_) = e^-*αTav*^ where *α* is the speed of the predator. Likewise, the probability of catching a prey, *C(T*_*ap*_), decreases with *T*_*ap*_ according to *C(T*_*ap*_) = e^*- λTap*^ where *λ* represents the speed of the prey. Note that when the individual approaches and catches prey, a quantity denoted by *m* is added to the corresponding value of *V*, reflecting the benefit of successfully capturing the prey.

With values for parameters *V, m, α*, and *λ*, it is possible to find the optimal thresholds for approaching and avoiding, which maximize the expected value for different values of *p* (probability of encountering a predator). We use *h* to denote the value of the approach threshold and *d* to denote the avoid threshold. The procedure for calculating the optimal thresholds is described in Appendix A.

### State-dependent drift diffusion model

DDMs usually assume that every time a decision is made, that decision is independent of any past or future experiences or circumstances (Trimmer et al., 2013; Nguyen et al., 2019; Gupta et al., 2024). However, because actions can alter the state of the individual, which in turn can influence future decisions, even when the environmental state is independent from one time step to another, the decisions are not independent of one another. Under such circumstances, the optimal behaviour can be calculated using state-dependent modelling (or dynamic programming) (Mangel & Clark, 1988; Houston & McNamara, 1999). This approach allows us to consider a sequence of decisions and allows the expected reproductive value, *V*, to emerge from the model rather than being a fixed input value (Houston & McNamara, 1999). Under this framework, the reproductive value is commonly approximated with a surrogate currency like expected reproductive success (i.e. number of offspring) using dynamic programming (Houston & McNamara, 1999).

A key feature of the state-dependent modelling framework is the incorporation of state variables (Houston & McNamara, 1999; Clark & Mangel, 2000). These variables typically represent an organism’s physiological state (e.g., energy reserves, body mass, parasite load) or environmental factors (e.g., temperature, prey density, geographical location) (Houston & McNamara, 1999). The importance of state variables lies in their ability to influence the effects of an organism’s actions on its survival and reproduction (Houston & McNamara, 1999). In our models, we will use energy reserves as the state variable, with the payoffs for each combination of response and circumstance depending on the individual’s reserve level.

Integrating a DDM into the state-dependent framework allows the response thresholds *h* and *d* to vary as a function of the individual’s reserve level. For example, an individual receiving a signal from another animal may reach threshold *d*, for an avoidance response, faster than threshold *h*, for an approach response, when reserve levels are high.

The process used to derive the thresholds and expected values for each reserve level, *i*, works backwards in time, repeatedly calculating optimal thresholds and expected reproductive values, refining estimates from what might be very crude approximations to stable values. By working sufficiently far back in time, this approach means that the reproductive values become independent of assumptions about values at the final time (Houston & McNamara, 1999). The values of *V* at neighbouring reserve levels indicated in the payoff matrix can then be used to obtain optimal thresholds at each reserve level. This means that the resulting values may depend on the values at reserve levels either one step higher or lower, or even further beyond these levels. We use the obtained thresholds to update the expected values at a given level and update the thresholds using the previously updated values. By iterating this process until both the values and thresholds stabilize, we identify the optimal strategy that maximizes reproductive value. In Appendix B we describe in more detail how this iterative process works, and how it makes use of the calculations described in Appendix A.

For the simple foraging scenario previously described, we discretize time into a series of independent periods, which are assumed to be significantly longer than the time taken (by the DDM) to reach a decision within each time step. In each time step the individual receives a signal from an animal that might be a predator or a prey, and the individual makes a decision (using their DDM). We assume that death (whether from predation or starvation) occurs at the end of a period. Here we focus on modelling using two proxies for reproductive value, expected reproductive success and expected survival time. For simplicity, in both proxies, the resulting payoffs will depend only on the payoffs at reserve levels one step higher or lower, without considering further levels. One unit of reserves is lost in each time step if the individual succeeds at avoiding a predator or if it misses a prey. One unit of reserves is obtained if a prey is captured. If the individual approaches a predator or its reserves drop to zero, it will die.

### Maximizing expected reproductive success

In the model for expected reproductive success we assume that individuals act to maximize their expected number of offspring. We assume asexual reproduction, and that the individual reproduces once it attains a level of reserves we denote as *L*. Thus, reserves are represented by an integer value, {0, 1, …, *L*}. We set *L* = 4 for all simulations to expedite calculations and demonstrate in Appendix C that similar qualitative results are obtained when using a higher number of reserve levels. At reserves of 1, the individual must approach the other animal, as it would die anyway at the end of the period if it avoids it. As a result, although the individual will not starve to death, the threat of starvation means that, at minimal reserves, the individual will accept any level of predation risk (McNamara, 1990).

When the individual reaches reserves of *L*, one unit is added to its payoff of reproductive success, but its reserves are reduced in the process by some amount we denote as *b*, which is the cost of producing one offspring. Therefore, when *L* is reached the payoff is *V(L – b) + 1*. Table 2 shows the payoff matrix for strategies that maximize reproductive success.

**Table 2.** Payoff matrix for the S3DM when maximizing reproductive success.

|  |  | Response |  |
| --- | --- | --- | --- |
|  |  | Approach | Avoid |
| Environmental condition | Predator | 0 | $S(T_{av}) V(i - 1)$ |
| | Prey | $C(T_{ap}) V(i + 1) + (1 - C(T_{ap})) V(i - 1)$ | $V(i - 1)$ |

Note that in contrast to the standard DDM, for the S3DM the benefit of successfully capturing the prey is reflected by a one unit increase in reserves.

### Maximizing expected survival time

When it comes to the model for maximizing expected survival time, there are some minor differences with respect to maximizing reproductive success. Here, we interpret each payoff value as representing an expected lifespan. We set a maximum limit for reserve levels, with reserves once again represented as integer values, {0, 1, …, *L*}. In a manner similar to reproductive success, when reserves are at 1, the individual must approach another animal, as avoiding it would result in death at the end of the period. At maximum reserves, *L*, if the individual captures a prey, it will maintain its reserve level. Table 3 shows the payoff matrix for strategies that maximize survival time.

**Table 3.** Payoff matrix for the S3DM when maximizing survival time.

|  |  | Approach | Avoid |
| --- | --- | --- | --- |
| Environmental condition | Predator | 1 | $S(T_{av}) V(i - 1) + 1$ |
| | Prey | $C(T_{ap})V(i + 1) + (1 - C(T_{ap}))V(i - 1) + 1$ | $V(i - 1) + 1$ |

## Results

To highlight the contrast between the outcomes of the two kinds of models we used the payoff values of the S3DM at an intermediate reserve level as input for the DDM and evaluate the expected value of gaining or losing a unit of food. Figure 1 shows a clear contrast between the thresholds for the two kinds of models across a range of *p* values when the goal is maximizing reproductive success. For the S3DM, as the probability of encountering a predator increases, individuals should spend more time (i.e., be slower but more accurate) deciding on an avoidance response and less time (i.e., be faster but less accurate) on an approach.

**Figure 1.**
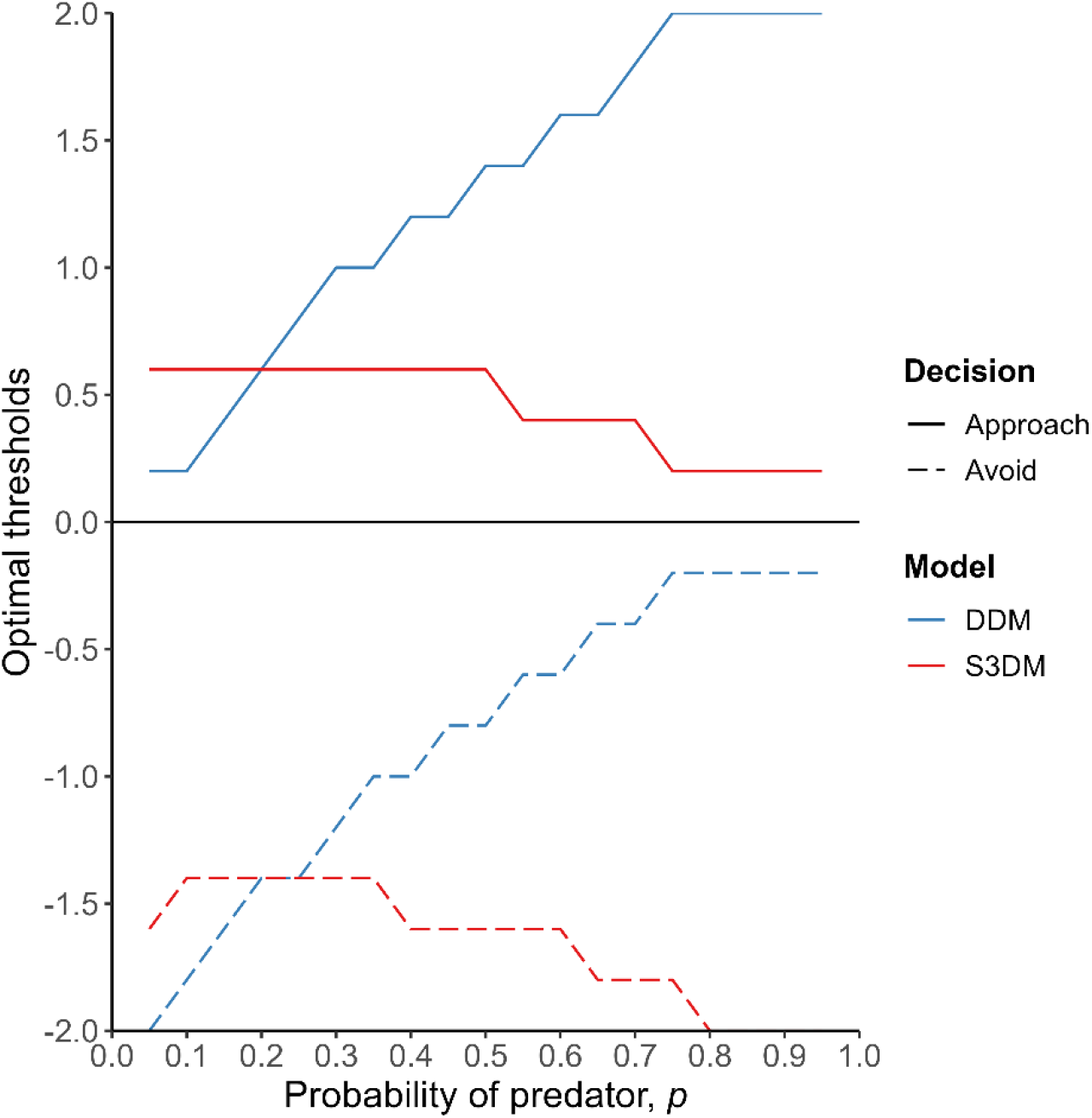
Optimal response thresholds for approach (solid lines) and avoidance (dashed lines) as a function of *p* for both the S3DM (red) and the DDM (blue) when maximizing reproductive success. We used the payoff values of the S3DM at a reserve level of 2 and *p* = 0.2 to draw the lines for the DDM (this provides payoff values of *V*(1) = 1.66 and *V*(3) = 2.58). Consequently, the lines coincide at that point. (Parameters: *α* = 0.2, *λ* = 0.2, *c* = 2, *L* = 4.)

However, the standard DDM predicts the opposite trend, with individuals expected to make avoidance decisions more quickly and take longer to decide on approach responses when encountering another animal.

These findings suggest that using the standard DDM to model decision-making in natural environments can lead to misleading conclusions due to its assumption that each decision is independent. When decisions are viewed in isolation, without considering state-dependent variables, it might seem logical that as the likelihood of encountering a predator increases, an individual should decide more quickly to avoid and take longer to approach, given the higher risk of predation. However, within a framework where multiple decisions impact an individual’s reserve level, the opposite behaviour may be more appropriate in certain circumstances.

From the S3DM perspective, when the goal is to accumulate reserves for reproduction, an individual should take more time to decide on avoidance when the risk of predation is high because this risk affects future decisions. If an individual quickly avoids an animal, as the standard DDM predicts, the risk may simply be deferred. A greater tendency to avoid predators increases the likelihood of depleting reserve levels, which can be detrimental when the goal is to maximize reserves for reproduction. In other words, when risks are high, the individual is more willing to be less accurate and risk making a mistake when approaching another animal, as it needs to reach the highest reserve level to reproduce.

The contrast between the two models is also evident in Figure 2. The standard DDM predicts a slower decline in expected reproductive success, suggesting that an individual might still reproduce under high predation risk, despite consistently avoiding and thus not catching prey. This is clearly unrealistic because animals need additional reserves to reproduce. The DDM’s assumption of isolated decisions that do not influence subsequent conditions or decisions can lead to these improbable predictions.

**Figure 2.**
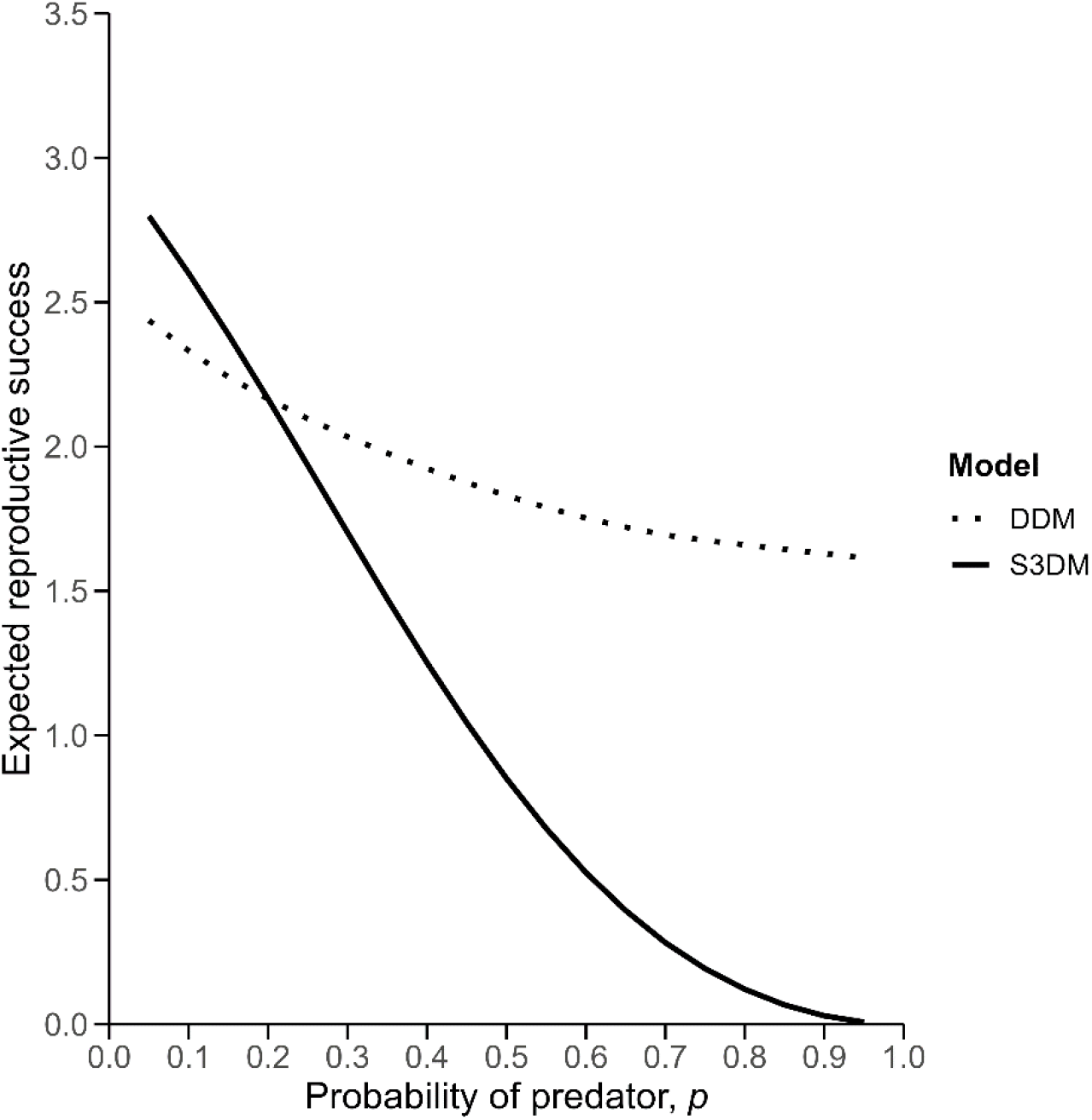
Expected reproductive success as a function of *p* for both the S3DM (solid line) and the DDM (dotted line). The payoff values of the S3DM at a reserve level of 2 and *p* = 0.2 were used to calculate the lines for the DDM (this provides payoff values of *V*(1) = 1.66 and *V*(3) = 2.58). Consequently, the lines coincide at that point. (Parameters: *α* = 0.2, *λ* = 0.2, *c* = 2, *L* = 4.)

The predictions of the standard DDM and the S3DMs are not always in opposite directions. Figure 3 illustrates the contrast between the outcomes of the two models when maximizing survival time. Once again, we used the payoff values of the S3DM at an intermediate reserve level as input for the DDM and evaluate the expected value of gaining or losing a unit of food. In this scenario, the S3DM predicts that individuals should take less time to decide on avoiding another animal and more time to decide on approaching it.

**Figure 3.**
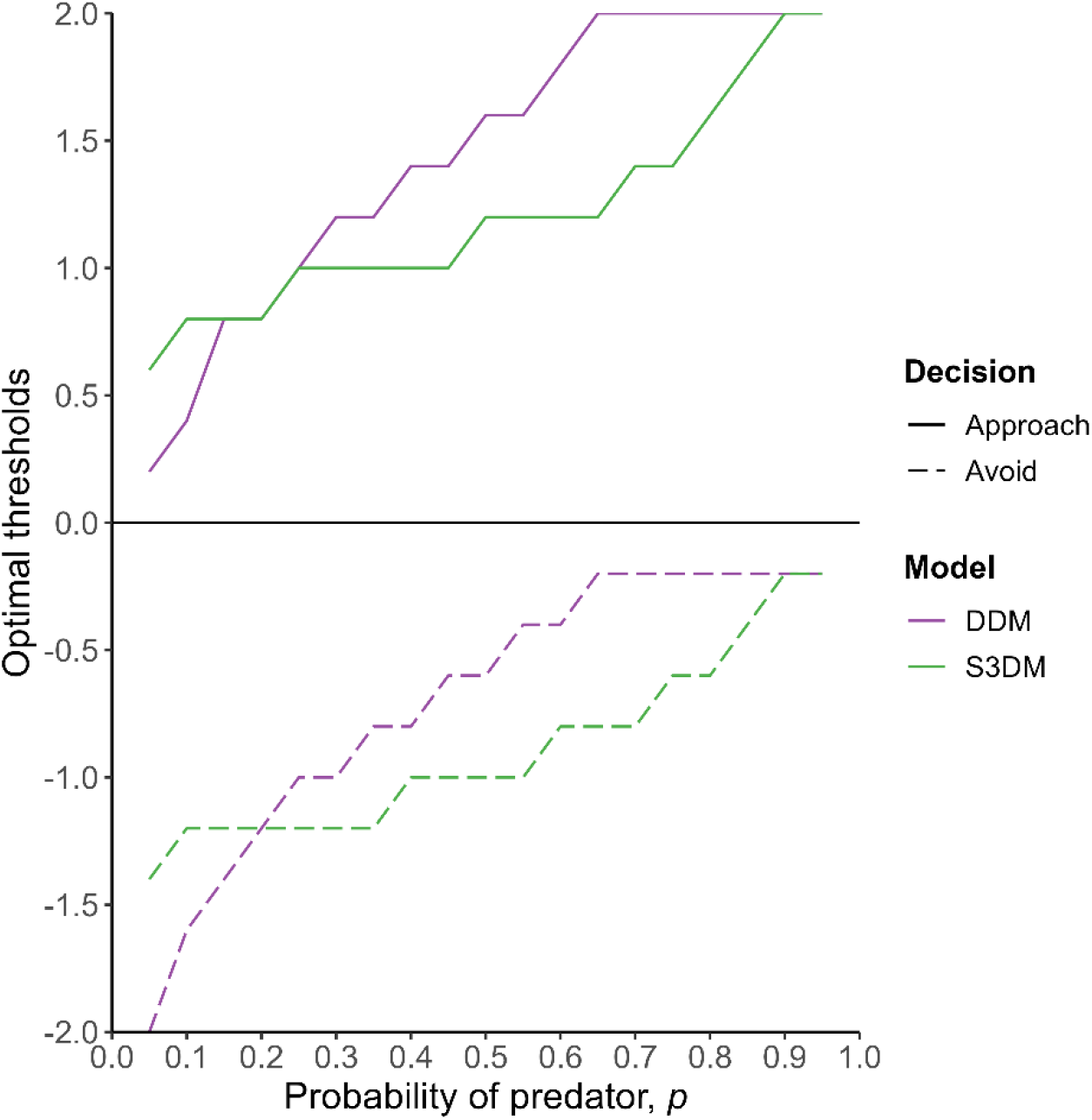
Optimal response thresholds for approach (solid lines) and avoidance (dashed lines) as a function of *p* for both the S3DM (green) and the DDM (purple) when maximizing survival time. We used the payoff values of the S3DM at a reserve level of 2 and *p* = 0.2 to draw the lines for the DDM (this provides payoff values of *V*(1) = 16.32 and *V*(3) = 21.85). Consequently, the lines coincide at that point. (Parameters: *α* = 0.2, *λ* = 0.2.)

The standard DDM also predicts increasing thresholds, but these thresholds start lower at small values of *p* and rise more quickly than those in the S3DM. As a result, at low levels of predation risk, the DDM suggests that individuals should make quicker decisions to approach another animal and slower decisions to avoid it compared to the S3DM. However, at medium and high levels of predation risk, this pattern reverses, with the DDM predicting slower decisions to approach and faster decisions to avoid than the S3DM. Overall, these results indicate that while the standard DDM and the S3DMs do not always produce entirely opposite trends, their predictions about how responses change with varying risk levels still differ noticeably.

The contrast between the DDM and the S3DM is more apparent in Figure 4. The S3DM predicts a rapid decline in expected survival time as predation risk increases. In comparison, the standard DDM predicts only a very small change in expected survival time, with a decrease of less than 5 units across the entire range of predator probabilities. This seems improbable because it suggests that the individual manages to survive for nearly the same amount of time, even when it is supposed to be constantly escaping under high predation risk.

**Figure 4.**
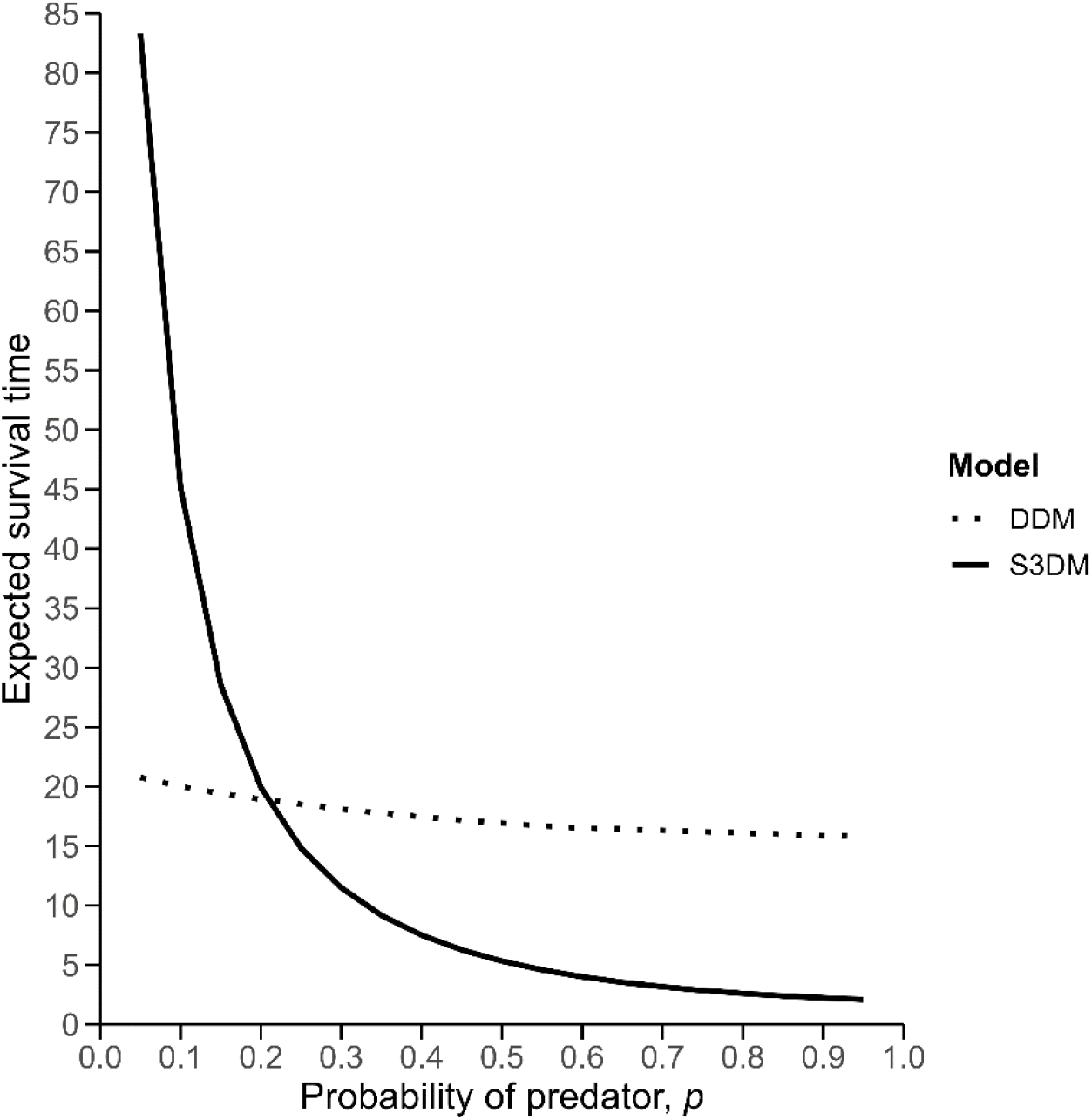
Expected survival time as a function of *p* for both the S3DM (solid line) and the DDM (dotted line). We used the payoff values of the S3DM at a reserve level of 2 and *p* = 0.2 to draw the lines for the DDM (this provides payoff values of *V*(1) = 16.32 and *V*(3) = 21.85). Consequently, the lines coincide at that point. (Parameters: *α* = 0.2, *λ* = 0.2.)

The differences in threshold trends between the S3DM for maximizing reproductive success and the S3DM for maximizing survival time can be explained as follows. If an individuals’ goal is to increase reserves to a reproductive level, then avoidance moves them further from their goal, and as predator prevalence, *p*, increases, the long-term prospects of overcoming that two-step difference reduces. In contrast, if the individual’s primary goal is maximising survival time, then each additional time step survived is adding to the value being achieved; it can therefore make more sense to avoid encountered animals as *p* increases.

## Discussion

Diffusion models have been extensively used to study simple two-choice decision processes and have successfully accounted for a substantial amount of behavioural data across various laboratory tasks and experimental conditions (Ratcliff & McKoon, 2008). More recently there has been a growing interest in the application of diffusion models to decision-making in more natural environments (Nguyen et al., 2019; Kilpatrick et al., 2019; Davidson & El Hady, 2019; Bidari et al., 2022). This work shows how DDMs can be placed in a state-dependent modelling framework, allowing analyses of sequential decisions (Nguyen et al., 2019; Houston et al., 2024; Houston & McNamara, 1982).

Considering a simple foraging scenario, we showed how behavioural predictions of the S3DMs may differ to those of a standard DDM. Specifically, in biological scenarios where natural selection favours organisms that maximize reproductive success, predictions of the standard DDM may be reversed. In such cases, if the chance of encountering a predator is increased, individuals should become less cautious and more quickly decide to approach another animal – which may seem counterintuitive. If the individual is more willing to mistakenly avoid another animal when predators are common, its reserves might continuously decrease and reduce its chances to reproduce. Congruent results have been found for maximization of reproductive success in the case of Signal Detection Theory (SDT) when incorporated into the state-dependent framework with the State-dependent Detection Theory (SDDT; Trimmer et al., 2017b).

Nevertheless, a standard DDM and the S3DMs do not always predict different trends. When natural selection favours individuals that maximize survival time, the trends can be in similar directions. Under this scenario, individuals should be more willing to mistakenly avoid another animal because reaching a high level of reserves is not critical for long-term survival. Even though state-dependent modelling could bring more realism to traditional models like SDT and DDMs, that does not necessarily mean that predictions made by these models will be completely reversed. Further research could compare the standard DDM predictions with those of the S3DMs using different proxies adequate for other biological scenarios (e.g., survival probability over a fixed period; Houston & McNamara, 1993).

S3DMs provide a clear integration of functional and mechanistic explanations of behaviour, consistent with Tinbergen’s conceptualization (Tinbergen, 1963; Font, 2023). The importance of integrating these forms of explanation has been highlighted by researchers in the fields of behavioural ecology and decision making (McNamara & Houston, 2009; Monaghan, 2014; Healy & Braithwaite, 2000). This integrative approach, termed “Evo-mecho”, acknowledges that purely functional accounts of behaviour might not be realistic, as they assume that behaviour depends on circumstances in a completely flexible way (McNamara & Houston, 2009; Fawcett et al., 2013; Fawcett et al., 2015; Giraldeau & Dubois, 2008). At the same time, it recognizes that research on the mechanisms of decision-making would benefit from considering what functions such mechanisms should fulfil (e.g., reproductive success) (McNamara & Houston, 2009; Bateson et al., 2011; Sherry, 2005). S3DMs include, as an essential component, the function of the mechanisms that underlie behaviour. Moreover, diffusion models, on which the S3DMs are based, have shown impressive correspondence with the neurological substrate of behaviour under some conditions (Brunton et al., 2013; Roitman & Shadlen, 2002; Bode et al., 2012; Polanía et al., 2014; Ratcliff et al., 2003, 2007; Smith & Ratcliff, 2004; Van Vugt et al., 2012; Ratcliff & McKoon, 2008), providing confidence that these models offer biologically plausible approximations. Thus, S3DMs may provide a valuable route forward for investigating the biological mechanisms underlying decision-making.

Our results also highlight the importance of carefully evaluating under which circumstances it is more reasonable to use single-decision models like the standard DDM over multiple-decision models. Single-decision models may suffice when modelling scenarios where an organism makes a crucial, once-in-a-lifetime decision (Trimmer et al., 2017b). For instance, species that show obligate monogamy and might have a unique opportunity to choose a mate (Kvarnemo, 2018). However, DDMs of decision-making are usually concerned with decisions that are made fast and considered at a “low cognitive level” (Ratcliff et al., 2016). For animals, most types of decisions are made repeatedly throughout their lives and for some these need to be made fast to survive. Thus, we consider that natural scenarios of foraging under risk are often better modelled using multiple-decision models like the S3DMs.

We have kept things simple with our foraging scenario; a promising avenue for future research is autocorrelation, where the probability of encountering a predator in one period partially predicts that probability in future periods. We have also assumed that an individual uses the same response thresholds during its whole life, so another avenue would be to consider how response thresholds should alter when learning processes increase discriminability of predators and prey.

Future research could identify scenarios where the predictions of the S3DMs diverge most significantly from standard DDMs. This would enable comparisons to be made using existing empirical findings, or guide new studies to determine whether and under what conditions S3DMs provide more accurate predictions of observed behaviours. If S3DMs demonstrate superior explanatory power across a broad range of contexts, this would underscore their potential as valuable tools for understanding decision-making processes.

