## Supplementary material for "Generalizing Drift Diffusion Models with a State-Dependent Framework"

### Overview

Here, we provide modelling details, proofs of the main text findings, and the code used. This document is divided into four sections:

**Appendix A: Calculating optimal thresholds in the standard DDM of decision-making**

This provides the general procedure required to calculate optimal thresholds for the models described in the main text. It also includes the code for the DDM that uses payoffs values of the S3DMs as input.

**Appendix B: Calculating thresholds and payoffs in the S3DMs**

This describes the system of dependencies for calculating optimal thresholds and payoffs for the S3DMs.

**Appendix C: Using a higher number of reserves levels**

This provides a proof that the results described in the main text are qualitatively the same when using a higher number of possible reserves levels.

### Appendix A: Calculating optimal thresholds in the standard DDM of decision-making

The decision-making time, *T*, may vary depending on whether the decision made is to approach or avoid an encountered animal. In this context, $T_{ap}$ represents the time taken to decide to approach, assuming that this decision is reached first, while $T_{av}$ represents the time taken to decide to avoid, assuming that this decision is reached first. We assume that the probability of escaping from a predator, $S(T_{av})$, decreases with time to reach the decision, $T_{av}$, according to $S(T_{av}) = e^{-\alpha T_{av}}$ where $\alpha$ is the speed of the predator. Likewise, the probability of catching a prey, $C(T_{ap})$, decreases with $T_{ap}$ according to $C(T_{ap}) = e^{-\lambda T_{ap}}$ where $\lambda$ represents the speed of the prey. When the individual approaches and catches prey, a quantity denoted by *m* is added to the corresponding value of *V*, reflecting the additional benefit of capturing the prey. These details are summarized in Table 1 of the main text.

The probability of the encountered animal being a predator is *p*, so the probability that it is a prey is (1 - *p*). The expected value for a given set of thresholds *h* and *d* is:

$$\mathrm{E}(\text{value}|\ d,h) =$$

$$p\ [\int_0^\infty (\text{P(reaching the approach threshold when it is a predator)}) \\ (\text{E(payoff) for approaching a predator at time t)}dt)$$

+

$$\int_0^\infty (\text{P(reaching the avoid threshold when it is a predator)}) \\ (\text{E(payoff) for avoiding a predator at time t)}dt)]$$

+

$$(1-p)[\int_0^\infty (\text{P(reaching the approach threshold when it is a prey)} \\ (\text{E(payoff) for approaching a prey at time t)}dt$$

+

$$\int_0^\infty (\text{P(reaching the avoid threshold when it is a prey)} \\ (\text{E(payoff) for avoiding a prey at time t)}dt)]$$

Consulting the handbook of Brownian motion [51], we can determine the four probabilities needed in the formula. We set $x = 0$ as the starting point of the drift diffusion process and $\mu = 1$ as the mean drift of information with time. We dispose that the drift is positive when the novel stimulus comes from a prey and negative when it comes from a predator. Thus, when we also include the expected payoffs shown in Table 1, the expected value for a given set of thresholds *h* and *d* is:

$$\mathrm{E}(\text{value}|\ d,h) =$$

$$p\int_0^\infty \mathrm{e}^{-\mu(h-x)-\frac{\mu^2 t}{2}} \sum_{k=-\infty}^{\infty} \frac{h-x+2k(h-d)}{\sqrt{2\pi}t^{3/2}} \mathrm{e}^{-\frac{(h-x+2k(h-d))^2}{2t}} V(\text{approach predator})\ dt$$

$$+p\int_0^\infty \mathrm{e}^{-\mu(d-x)-\frac{\mu^2 t}{2}}\sum_{k=-\infty}^{\infty}\frac{x-d+2k(h-d)}{\sqrt{2\pi}t^{3/2}}\mathrm{e}^{-\frac{(x-d+2k(h-d))^2}{2t}}Ve^{-\alpha T_{av}}\,dt$$

$$+(1-p)\int_0^\infty \mathrm{e}^{\mu(h-x)-\frac{\mu^2 t}{2}}\sum_{k=-\infty}^{\infty}\frac{h-x+2k(h-d)}{\sqrt{2\pi}t^{\frac{3}{2}}}\mathrm{e}^{-\frac{(h-x+2k(h-d))^2}{2t}}((V+m)e^{-\lambda T_{ap}}+V(1-e^{-\lambda T_{ap}}))\,dt$$

$$+(1-p)\int_0^\infty \mathrm{e}^{\mu(d-x)-\frac{\mu^2 t}{2}}\sum_{k=-\infty}^{\infty}\frac{x-d+2k(h-d)}{\sqrt{2\pi}t^{3/2}}\mathrm{e}^{-\frac{(x-d+2k(h-d))^2}{2t}}V\,dt$$

As displayed in Table 1 of the main text, we are setting the payoff for approaching a predator to 0. We approximate the sum over the infinite *k* sum by running between finite limits of [-10, 10]. We approximate the infinite time integral by just running up to a time of 10 seconds, when we consider a decision is pretty much bound to be made, by numerically summing time slices of 0.1 seconds. We perform the integration using the Gauss-Kronrod method. In order to obtain more precise results, we suggest integrating over a time of 100 seconds or more with time slices of 0.001 seconds; values that we could not use due to time constraints of the project and the need of more computational power.

By determining the values of *d* and *h* that maximize E(value|*d,h*), we can identify the optimal thresholds that align with the specific environment (i.e., given *p, μ, λ, α, V,* and *m*).

The following code was used to calculate the DDM values for Figure 1 and Figure 2 in the main text. In this code, the payoffs are replaced with the payoff values of the S3DM for maximizing reproductive success at a reserve level of 2 and *p* = 0.2. The code is as follows:

```
library(pracma)

library(ggplot2)

library(dplyr)

library(ggtext)


# Define constants

alpha <- 0.2 # Speed of predator

lambda <- 0.2 # Speed of prey

mu <- 1 # Mean drift of information with time

x <- 0 # Starting point

time_step <- 0.1 # Time step for numerical integration

time_limit <- 10 # Time limit for numerical integration

k_limit <- 10  # Approximate the sum over the infinite k sum by running between limits of [-10, 10]
```

```
# Integrands
# Integrand for approaching a predator
integrand_ap_pr <- function(t, d, h) {
  sum_k <- 0
  for (k in -k_limit:k_limit) {
    term <- ((h - x + 2 * k * (h - d)) / (sqrt(2 * pi) * t^(3/2))) * exp(-(h - x + 2 * k * (h - d))^2 / (2 * t))
    sum_k <- sum_k + term
  }
  result <- exp(-mu * (h - x) - ((mu^2) * t / 2)) * sum_k * 0
  return(result)
}
# Integrand for avoiding a predator
integrand_av_pr <- function(t, d, h) {
  sum_k <- 0
  for (k in -k_limit:k_limit) {
    term <- ((x - d + 2 * k * (h - d)) / (sqrt(2 * pi) * t^(3/2))) * exp(-(x - d + 2 * k * (h - d))^2 / (2 * t))
    sum_k <- sum_k + term
  }
  result <- exp(-mu * (d - x) - ((mu^2) * t / 2)) * sum_k * (1.664047 * exp(-alpha * t))
  return(result)
}
# Integrand for approaching a prey
integrand_ap_py <- function(t, d, h) {
  sum_k <- 0
  for (k in -k_limit:k_limit) {
    term <- ((h - x + 2 * k * (h - d)) / (sqrt(2 * pi) * t^(3/2))) * exp(-(h - x + 2 * k * (h - d))^2 / (2 * t))
    sum_k <- sum_k + term
  }
```

```
  result <- exp(mu * (h - x) - ((mu^2) * t / 2)) * sum_k * ((2.577832 * exp(-lambda * t)) + (1.664047
* (1 - exp(-lambda * t))))
  return(result)
}
# Integrand for avoiding a prey
integrand_av_py <- function(t, d, h) {
  sum_k <- 0
  for (k in -k_limit:k_limit) {
    term <- ((x - d + 2 * k * (h - d)) / (sqrt(2 * pi) * t^(3/2))) * exp(-(x - d + 2 * k * (h - d))^2 / (2 * t))
    sum_k <- sum_k + term
  }
  result <- exp(mu * (d - x) - ((mu^2) * t / 2)) * sum_k * 1.664047
  return(result)
}

# Calculate the expected reward given thresholds d and h
expected_reward <- function(d, h, p) {
  E_ap_pr <- pracma::integral(integrand_ap_pr, xmin = 0, xmax = time_limit,
                              d = d, h = h, method = "Kronrod",
                              random = F, no_intervals = time_limit/time_step)
  E_av_pr <- pracma::integral(integrand_av_pr, xmin = 0, xmax = time_limit,
                              d = d, h = h, method = "Kronrod",
                              random = F, no_intervals = time_limit/time_step)
  E_ap_py <- pracma::integral(integrand_ap_py, xmin = 0, xmax = time_limit,
                              d = d, h = h, method = "Kronrod",
                              random = F, no_intervals = time_limit/time_step)
  E_av_py <- pracma::integral(integrand_av_py, xmin = 0, xmax = time_limit,
                              d = d, h = h, method = "Kronrod",
                              random = F, no_intervals = time_limit/time_step)
```

```
  E_reward <- p * E_ap_pr + p * E_av_pr + (1 - p) * E_ap_py + (1 - p) * E_av_py
  return(E_reward)
}

# Optimize the thresholds d and h to maximize the expected reward
optimize_thresholds <- function(p) {
  d_range <- seq(-0.2, -2, by = -0.2)
  h_range <- seq(0.2, 2, by = 0.2)
  max_reward <- -Inf
  th_d <- NA
  th_h <- NA

  for (d in d_range) {
    for (h in h_range) {
      reward <- expected_reward(d, h, p)
      if (reward > max_reward) {
        max_reward <- reward
        th_d <- d
        th_h <- h
      }
    }
  }

  return(list(th_h = th_h, th_d = th_d, max_reward = max_reward))
}

# Run the optimization for different values of p
p_values <- seq(0.05, 0.95, 0.05)
```

```
results_3 <- data.frame(p = numeric(), th_h = numeric(), th_d = numeric(), max_reward =
numeric())


for (p in p_values) {
  res <- optimize_thresholds(p)
  results_3 <- rbind(results_3, data.frame(p = p, th_h = res$th_h, th_d = res$th_d, max_reward =
res$max_reward))
}


# Plot Optimal thresholds vs Probability of predator
p3 <- ggplot(results_3, aes(x = p)) +
  geom_line(aes(y = th_h, color = "Approach")) +
  geom_line(aes(y = th_d, color = "Avoid")) +
  scale_x_continuous(name = "Probability of predator, *p*",
                     limits = c(0, 1), breaks = seq(0, 1, 0.1)) +
  scale_y_continuous(name = "Optimal thresholds",
                     limits = c(-2, 2), breaks = seq(-2, 2, 0.5)) +
  scale_color_manual(values = c("Avoid" = "blue", "Approach" = "red")) +
  theme(legend.position = "none") +
  annotate("text", x = 0.4, y = 1.5, label = "Approach", color = "red") +
  annotate("text", x = 0.4, y = -1.5, label = "Avoid", color = "blue") +
  theme_classic() +
  theme(axis.title.x = ggtext::element_markdown(), legend.position = "none")


# Plot Expected reproductive success vs Probability of predator
p7 <- ggplot(results_3, aes(x = p, y = max_reward)) +
  geom_line(color = "black", linewidth = 1) +
  geom_point(shape = 10, size = 3, color = "black") +
  scale_x_continuous(name = "Probability of predator, *p*",
                     limits = c(0, 1),
```

```
                  breaks = seq(0, 1, 0.1),
                  expand = c(0, 0)) +
  scale_y_continuous(name = "Expected reproductive success",
                  limits = c(0, 3.5),
                  breaks = c(0, 0.5, 1.0, 1.5, 2.0, 2.5, 3.0, 3.5),
                  expand = c(0, 0)) +
  theme_classic() +
  theme(axis.title.x = ggtext::element_markdown(size = 12),
        axis.title.y = element_text(size = 12),
        axis.line.x = element_line(color = "black"),
        axis.line.y = element_line(color = "black"),
        panel.grid = element_blank(),
        axis.ticks = element_line(color = "black"),
        axis.ticks.length = unit(0.2, "cm"),
        legend.position = "none")
```

The following code was used to calculate the DDM values for Figure 3 and Figure 4 in the main text. In this code, the payoffs are replaced with the payoff values of the S3DM for maximizing survival time at a reserve level of 2 and *p* = 0.2. The code is as follows:

```
library(pracma)
library(ggplot2)
library(dplyr)
library(ggtext)


# Define constants
alpha <- 0.2 # Speed of predator
lambda <- 0.2 # Speed of prey
mu <- 1 # Mean drift of information with time
x <- 0 # Starting point
time_step <- 0.1 # Time step for numerical integration
```

```
time_limit <- 10 # Time limit for numerical integration
k_limit <- 10  # Approximate the sum over the infinite k sum by running between limits of [-10, 10]

# Integrands

# Integrand for approaching a predator
integrand_ap_pr <- function(t, d, h) {
  sum_k <- 0
  for (k in -k_limit:k_limit) {
    term <- ((h - x + 2 * k * (h - d)) / (sqrt(2 * pi) * t^(3/2))) * exp(-(h - x + 2 * k * (h - d))^2 / (2 * t))
    sum_k <- sum_k + term
  }
  result <- exp(-mu * (h - x) - ((mu^2) * t / 2)) * sum_k * 0
  return(result)
}
# Integrand for avoiding a predator
integrand_av_pr <- function(t, d, h) {
  sum_k <- 0
  for (k in -k_limit:k_limit) {
    term <- ((x - d + 2 * k * (h - d)) / (sqrt(2 * pi) * t^(3/2))) * exp(-(x - d + 2 * k * (h - d))^2 / (2 * t))
    sum_k <- sum_k + term
  }
  result <- exp(-mu * (d - x) - ((mu^2) * t / 2)) * sum_k * ((16.31694* exp(-alpha * t)))
  return(result)
}
# Integrand for approaching a prey
integrand_ap_py <- function(t, d, h) {
  sum_k <- 0
  for (k in -k_limit:k_limit) {
```

```
    term <- ((h - x + 2 * k * (h - d)) / (sqrt(2 * pi) * t^(3/2))) * exp(-(h - x + 2 * k * (h - d))^2 / (2 * t))
    sum_k <- sum_k + term
  }
  result <- exp(mu * (h - x) - ((mu^2) * t / 2)) * sum_k * ((21.84652 * exp(-lambda * t)) + (16.31694
* (1 - exp(-lambda * t))))
  return(result)
}
# Integrand for avoiding a prey
integrand_av_py <- function(t, d, h) {
  sum_k <- 0
  for (k in -k_limit:k_limit) {
    term <- ((x - d + 2 * k * (h - d)) / (sqrt(2 * pi) * t^(3/2))) * exp(-(x - d + 2 * k * (h - d))^2 / (2 * t))
    sum_k <- sum_k + term
  }
  result <- exp(mu * (d - x) - ((mu^2) * t / 2)) * sum_k * 16.31694
  return(result)
}


# Calculate the expected reward given thresholds d and h
expected_reward <- function(d, h, p) {
  E_ap_pr <- pracma::integral(integrand_ap_pr, xmin = 0, xmax = time_limit,
                              d = d, h = h, method = "Kronrod",
                              random = F, no_intervals = time_limit/time_step)
  E_av_pr <- pracma::integral(integrand_av_pr, xmin = 0, xmax = time_limit,
                              d = d, h = h, method = "Kronrod",
                              random = F, no_intervals = time_limit/time_step)
  E_ap_py <- pracma::integral(integrand_ap_py, xmin = 0, xmax = time_limit,
                              d = d, h = h, method = "Kronrod",
                              random = F, no_intervals = time_limit/time_step)
```

```
  E_av_py <- pracma::integral(integrand_av_py, xmin = 0, xmax = time_limit,
                              d = d, h = h, method = "Kronrod",
                              random = F, no_intervals = time_limit/time_step)

  E_reward <- p * E_ap_pr + p * E_av_pr + (1 - p) * E_ap_py + (1 - p) * E_av_py
  return(E_reward)
}

# Optimize the thresholds d and h to maximize the expected reward
optimize_thresholds <- function(p) {
  d_range <- seq(-0.2, -2, by = -0.2)
  h_range <- seq(0.2, 2, by = 0.2)
  max_reward <- -Inf
  th_d <- NA
  th_h <- NA

  for (d in d_range) {
    for (h in h_range) {
      reward <- expected_reward(d, h, p)
      if (reward > max_reward) {
        max_reward <- reward
        th_d <- d
        th_h <- h
      }
    }
  }

  return(list(th_h = th_h, th_d = th_d, max_reward = max_reward))
}
```

```
# Run the optimization for different values of p

p_values <- seq(0.05, 0.95, 0.05)

results_4 <- data.frame(p = numeric(), th_h = numeric(), th_d = numeric(), max_reward = numeric())


for (p in p_values) {

  res <- optimize_thresholds(p)

  results_4 <- rbind(results_4, data.frame(p = p, th_h = res$th_h, th_d = res$th_d, max_reward = res$max_reward))

}


# Plot Optimal thresholds vs Probability of predator

p4 <- ggplot(results_4, aes(x = p)) +

  geom_line(aes(y = th_h, color = "Approach")) +

  geom_line(aes(y = th_d, color = "Avoid")) +

  scale_x_continuous(name = "Probability of predator, *p*",

                     limits = c(0, 1), breaks = seq(0, 1, 0.1)) +

  scale_y_continuous(name = "Optimal thresholds",

                     limits = c(-2, 2), breaks = seq(-2, 2, 0.5)) +

  scale_color_manual(values = c("Avoid" = "blue", "Approach" = "red")) +

  theme(legend.position = "none") +

  annotate("text", x = 0.4, y = 1.8, label = "Approach", color = "red") +

  annotate("text", x = 0.4, y = -1.1, label = "Avoid", color = "blue") +

  theme_classic() +

  theme(axis.title.x = ggtext::element_markdown(), legend.position = "none")


# Plot Expected survival time vs Probability of predator

p8 <- ggplot(results_4, aes(x = p, y = max_reward)) +

  geom_line(color = "black", linewidth = 1) +
```

```
geom_point(shape = 5, size = 3, color = "black") +
scale_x_continuous(name = "Probability of predator, *p*",
            limits = c(0, 1),
            breaks = seq(0, 1, 0.1),
            expand = c(0, 0)) +
scale_y_continuous(name = "Expected survival time",
            limits = c(0, 65),
            breaks = seq(0, 65, by = 5),
            expand = c(0, 0)) +
theme_classic() +
theme(axis.title.x = ggtext::element_markdown(size = 12),
    axis.title.y = element_text(size = 12),
    axis.line.x = element_line(color = "black"),
    axis.line.y = element_line(color = "black"),
    panel.grid = element_blank(),
    axis.ticks = element_line(color = "black"),
    axis.ticks.length = unit(0.2, "cm"),
    legend.position = "none")
```

**Appendix B: Calculating thresholds and payoffs in the S3DMs**

The process to calculate thresholds and payoffs in the S3DMs makes use of the same structure used in the E(value|*d,h*) formula shown in Appendix A. The changes to this formula are due to the dependence of the payoffs in Table 2 on the expected value at reserve levels either one step higher or lower, which we represent as *V(i + 1)* and *V(i - 1)*. These values at neighbouring reserves levels are used in the new formula. Thus, when maximizing reproductive success, the formula to calculate the expected value is:

$$\mathrm{E}(\text{value}|\ d,h) =$$

$$p\int_0^\infty \mathrm{e}^{-\mu(h-x)-\frac{\mu^2 t}{2}} \sum_{k=-\infty}^{\infty} \frac{h-x+2k(h-d)}{\sqrt{2\pi}t^{3/2}}\,\mathrm{e}^{-\frac{\left(h-x+2k(h-d)\right)^2}{2t}} V(\text{approach predator})dt$$

$$+\,p\int_0^\infty \mathrm{e}^{-\mu(d-x)-\frac{\mu^2 t}{2}} \sum_{k=-\infty}^{\infty} \frac{x-d+2k(h-d)}{\sqrt{2\pi}t^{\frac{3}{2}}}\mathrm{e}^{-\frac{\left(x-d+2k(h-d)\right)^2}{2t}} V(i-1)e^{-\alpha T_{av}}\,dt$$

$+ (1-p)\int_0^{\infty} \mathrm{e}^{\mu(h-x)-\frac{\mu^2 t}{2}} \sum_{k=-\infty}^{\infty} \frac{h-x+2k(h-d)}{\sqrt{2\pi} t^{\frac{3}{2}}} \mathrm{e}^{-\frac{(h-x+2k(h-d))^2}{2t}} (V(i+1)e^{-\lambda T_{ap}} + V(i-1)(1-e^{-\lambda T_{ap}}))\, dt$

$$+(1-p)\int_0^{\infty} \mathrm{e}^{\mu(d-x)-\frac{\mu^2 t}{2}} \sum_{k=-\infty}^{\infty} \frac{x-d+2k(h-d)}{\sqrt{2\pi} t^{\frac{3}{2}}} \mathrm{e}^{-\frac{(x-d+2k(h-d))^2}{2t}} V(i-1)\, dt$$

By determining the values of *d* and *h* that maximize E(value|*d,h*) we can identify the optimal thresholds at each reserves level that align with the specific environment (i.e., given *p, μ, λ, α,* and *V*). Using dynamic programming we can take the calculated expected values at neighbouring reserve levels to update the payoffs in Table 2 and then find another set of optimal thresholds. We ran this iterative process for 100 discrete time units, at which point the expected values had stabilized.

The following code was used to calculate the S3DM values for Figure 1 and Figure 2 in the main text. The code is as follows:

```
library(pracma)

library(ggplot2)

library(dplyr)

library(ggtext)


# Define constants

alpha <- 0.2 # Speed of predator

lambda <- 0.2 # Speed of prey

mu <- 1 # Mean drift of information with time

x <- 0 # Starting point

time_step <- 0.1 # Time step for numerical integration

time_limit <- 10 # Time limit for numerical integration

k_limit <- 10  # Approximate the sum over the infinite k sum by running between limits of [-10, 10]


# Create initial reserves and t columns

reserves <- c(4:0)

b_t <- c(4:0)
```

```
th_d <- rep(NA, 5)
th_h <- rep(NA, 5)
L <- 4
c <- 2

# Create initial dataframe
values <- data.frame(reserves, b_t)
thresholds <- data.frame(reserves, th_d, th_h)

# Integrands

# Integrand for approaching a predator
integrand_ap_pr <- function(t, d, h) {
  sum_k <- 0
  for (k in -k_limit:k_limit) {
    term <- ((h - x + 2 * k * (h - d)) / (sqrt(2 * pi) * t^(3/2))) * exp(-(h - x + 2 * k * (h - d))^2 / (2 * t))
    sum_k <- sum_k + term
  }
  result <- exp(-mu * (h - x) - ((mu^2) * t / 2)) * sum_k * 0
  return(result)
}
# Integrand for avoiding a predator
integrand_av_pr <- function(t, d, h, below_b_t) {
  sum_k <- 0
  for (k in -k_limit:k_limit) {
    term <- ((x - d + 2 * k * (h - d)) / (sqrt(2 * pi) * t^(3/2))) * exp(-(x - d + 2 * k * (h - d))^2 / (2 * t))
    sum_k <- sum_k + term
  }
```

```
  result <- exp(-mu * (d - x) - ((mu^2) * t / 2)) * sum_k * ((below_b_t * exp(-alpha * t)) + (0 * (1 -
exp(-alpha * t))))
  return(result)
}
# Integrand for approaching a prey
integrand_ap_py <- function(t, d, h, below_b_t, above_b_t) {
  sum_k <- 0
  for (k in -k_limit:k_limit) {
    term <- ((h - x + 2 * k * (h - d)) / (sqrt(2 * pi) * t^(3/2))) * exp(-(h - x + 2 * k * (h - d))^2 / (2 * t))
    sum_k <- sum_k + term
  }
  result <- exp(mu * (h - x) - ((mu^2) * t / 2)) * sum_k * ((above_b_t * exp(-lambda * t)) +
(below_b_t * (1 - exp(-lambda * t))))
  return(result)
}
# Integrand for avoiding a prey
integrand_av_py <- function(t, d, h, below_b_t) {
  sum_k <- 0
  for (k in -k_limit:k_limit) {
    term <- ((x - d + 2 * k * (h - d)) / (sqrt(2 * pi) * t^(3/2))) * exp(-(x - d + 2 * k * (h - d))^2 / (2 * t))
    sum_k <- sum_k + term
  }
  result <- exp(mu * (d - x) - ((mu^2) * t / 2)) * sum_k * below_b_t
  return(result)
}


# Calculate the expected reward given thresholds d and h
expected_reward <- function(d, h, p, below_b_t, above_b_t) {
  E_ap_pr <- pracma::integral(integrand_ap_pr, xmin = 0, xmax = time_limit,
                              d = d, h = h, method = "Kronrod",
```

```
                    random = F, no_intervals = time_limit/time_step)
  E_av_pr <- pracma::integral(integrand_av_pr, xmin = 0, xmax = time_limit,
                    d = d, h = h, below_b_t = below_b_t, method = "Kronrod",
                    random = F, no_intervals = time_limit/time_step)
  E_ap_py <- pracma::integral(integrand_ap_py, xmin = 0, xmax = time_limit,
                    d = d, h = h, below_b_t = below_b_t,
                    above_b_t = above_b_t, method = "Kronrod",
                    random = F, no_intervals = time_limit/time_step)
  E_av_py <- pracma::integral(integrand_av_py, xmin = 0, xmax = time_limit,
                    d = d, h = h, below_b_t = below_b_t, method = "Kronrod",
                    random = F, no_intervals = time_limit/time_step)

  E_reward <- p * E_ap_pr + p * E_av_pr + (1 - p) * E_ap_py + (1 - p) * E_av_py
  return(E_reward)
}

# Optimize the thresholds d and h to maximize the expected reward
optimize_thresholds <- function(p, below_b_t, above_b_t) {
  d_range <- seq(-0.2, -2, by = -0.2)
  h_range <- seq(0.2, 2, by = 0.2)
  max_reward <- -Inf
  optimal_d <- NA
  optimal_h <- NA

  for (d in d_range) {
    for (h in h_range) {
      reward <- expected_reward(d, h, p, below_b_t, above_b_t)
      if (reward > max_reward) {
        max_reward <- reward
```

```
        optimal_d <- d
        optimal_h <- h
      }
    }
  }

  return(list(optimal_h = optimal_h, optimal_d = optimal_d, max_reward = max_reward))
}

# Function to run the optimization and store results of 100 iterations
run_optimization <- function(p) {
  values <- data.frame(reserves, b_t)
  thresholds <- data.frame(reserves, th_d, th_h)

  for (i in 1:100) {
    new_b_t_col_name <- paste0("b_t-", i)
    new_th_d_col_name <- paste0("th_d-", i)
    new_th_h_col_name <- paste0("th_h-", i)

    values[[new_b_t_col_name]] <- NA
    thresholds[[new_th_d_col_name]] <- NA
    thresholds[[new_th_h_col_name]] <- NA

    prev_b_t_col_name <- if (i == 1) "b_t" else paste0("b_t-", i - 1)

    for (j in 1:nrow(values)) {
      if (values$reserves[j] == 0) {
        values[[new_b_t_col_name]][j] <- 0
        thresholds[[new_th_d_col_name]][j] <- NA
```

```
        thresholds[[new_th_h_col_name]][j] <- NA
      } else if (values$reserves[j] == L){
        values[[new_b_t_col_name]][j] <- (L - c) + 1
      } else {
        above_b_t <- if (j == 1) values[[prev_b_t_col_name]][j] else values[[prev_b_t_col_name]][j - 1]
        below_b_t <- if (j == nrow(values)) values[[prev_b_t_col_name]][j] else
values[[prev_b_t_col_name]][j + 1]

        list_thresholds_reward <- optimize_thresholds(p, below_b_t, above_b_t)
        thresholds[[new_th_d_col_name]][j] <- list_thresholds_reward$optimal_d
        thresholds[[new_th_h_col_name]][j] <- list_thresholds_reward$optimal_h
        values[[new_b_t_col_name]][j] <- list_thresholds_reward$max_reward
      }
    }
  }

  return(list(values = values, thresholds = thresholds))
}

# Run the optimization for different values of p
p_values <- seq(0.05, 0.95, 0.05)
results_list <- list()

for (p in p_values) {
  results <- run_optimization(p)
  results_list[[paste0("p_", p, "_values")]] <- results$values
  results_list[[paste0("p_", p, "_thresholds")]] <- results$thresholds
}
```

```
# Extract the thresholds for reserves = 2 of the 100th iteration
plot_data <- data.frame(p = p_values, th_d = NA, th_h = NA)

for (i in 1:length(p_values)) {
  p <- p_values[i]
  threshold_df <- results_list[[paste0("p_", p, "_thresholds")]]
  plot_data$th_d[i] <- threshold_df[3, "th_d-100"]
  plot_data$th_h[i] <- threshold_df[3, "th_h-100"]
}

# Plot Optimal thresholds vs Probability of predator
p1 <- ggplot(plot_data, aes(x = p)) +
  geom_line(aes(y = th_h, color = "Approach")) +
  geom_line(aes(y = th_d, color = "Avoid")) +
  labs(x = "probability of predator, *p*", y = "optimal thresholds") +
  scale_x_continuous(limits = c(0, 1), breaks = seq(0, 1, 0.1), expand = c(0, 0)) +
  scale_y_continuous(limits = c(-2, 2), breaks = seq(-2, 2, 0.5), expand = c(0, 0)) +
  scale_color_manual(values = c("Avoid" = "blue", "Approach" = "red")) +
  annotate("text", x = 0.4, y = 1, label = "Approach", color = "red") +
  annotate("text", x = 0.4, y = -1.8, label = "Avoid", color = "blue") +
  theme_classic() +
  theme(axis.title.x = ggtext::element_markdown(), legend.position = "none")

# Extract the expected values for reserves = 2 of the 100th iteration
plot_data_5 <- data.frame(p = p_values, exp_re_suc = NA)

for (i in 1:length(p_values)) {
  p <- p_values[i]
  exp_re_suc_df <- results_list[[paste0("p_", p, "_values")]]
```

```
  plot_data_5$exp_re_suc[i] <- exp_re_suc_df[3, "b_t-100"]
}


# Plot Expected reproductive success vs Probability of predator
p5 <- ggplot(plot_data_5, aes(x = p, y = exp_re_suc)) +
  geom_line(color = "black", linewidth = 1) +
  geom_point(shape = 8, size = 3, color = "black") +
  scale_x_continuous(name = "Probability of predator, *p*",
                     limits = c(0, 1),
                     breaks = seq(0, 1, 0.1),
                     expand = c(0, 0)) +
  scale_y_continuous(name = "Expected reproductive success",
                     limits = c(0, 3.5),
                     breaks = c(0, 0.5, 1.0, 1.5, 2.0, 2.5, 3.0, 3.5),
                     expand = c(0, 0)) +
  theme_classic() +
  theme(axis.title.x = ggtext::element_markdown(size = 12),
        axis.title.y = element_text(size = 12),
        axis.line.x = element_line(color = "black"),
        axis.line.y = element_line(color = "black"),
        panel.grid = element_blank(),
        axis.ticks = element_line(color = "black"),
        axis.ticks.length = unit(0.2, "cm"),
        legend.position = "none")
```

The process is the same when maximizing survival time, but this time we use the payoffs corresponding to those in Table 3 of the main text. Thus, when maximizing survival time, the formula to calculate the expected value is:

$\mathrm{E(value} |\ d,h) =$

$$p\int_0^{\infty} \mathrm{e}^{-\mu(h-x)-\frac{\mu^2 t}{2}} \sum_{k=-\infty}^{\infty} \frac{h-x+2k(h-d)}{\sqrt{2\pi}t^{3/2}} \mathrm{e}^{-\frac{\left(h-x+2k(h-d)\right)^2}{2t}} V(\text{approach predator}) + 1\, dt$$

$$+\,p\int_0^{\infty} \mathrm{e}^{-\mu(d-x)-\frac{\mu^2 t}{2}} \sum_{k=-\infty}^{\infty} \frac{x-d+2k(h-d)}{\sqrt{2\pi}t^{\frac{3}{2}}} \mathrm{e}^{-\frac{\left(x-d+2k(h-d)\right)^2}{2t}} (V(i-1)e^{-\alpha T_{av}} + 1)\, dt$$

$$+\,(1-p)\int_0^{\infty} \mathrm{e}^{\mu(h-x)-\frac{\mu^2 t}{2}} \sum_{k=-\infty}^{\infty} \frac{h-x+2k(h-d)}{\sqrt{2\pi}t^{\frac{3}{2}}} \mathrm{e}^{-\frac{\left(h-x+2k(h-d)\right)^2}{2t}} (V(i+1)e^{-\lambda T_{ap}} + (V(i-1)(1-e^{-\lambda T_{ap}}) + 1)\, dt$$

$$+(1-p)\int_0^{\infty} \mathrm{e}^{\mu(d-x)-\frac{\mu^2 t}{2}} \sum_{k=-\infty}^{\infty} \frac{x-d+2k(h-d)}{\sqrt{2\pi}t^{\frac{3}{2}}} \mathrm{e}^{-\frac{\left(x-d+2k(h-d)\right)^2}{2t}} (V(i-1)+1)\, dt$$

We ran the dynamic programming iterative process for 200 discrete time units, at which point the expected values had stabilized.

The following code was used to calculate the S3DM values for Figure 3 and Figure 4 in the main text. The code is as follows:

```
library(pracma)

library(ggplot2)

library(dplyr)

library(ggtext)


# Define constants

alpha <- 0.2 # Speed of predator

lambda <- 0.2 # Speed of prey

mu <- 1 # Mean drift of information with time

x <- 0 # Starting point

time_step <- 0.1 # Time step for numerical integration

time_limit <- 10 # Time limit for numerical integration

k_limit <- 10 # Approximate the sum over the infinite k sum by running between limits of [-10, 10]


# Create initial reserves and t columns

reserves <- c(4:0)
```

```
b_t <- c(4:0)
th_d <- rep(NA, 5)
th_h <- rep(NA, 5)

# Create initial dataframe
values <- data.frame(reserves, b_t)
thresholds <- data.frame(reserves, th_d, th_h)

# Integrands

# Integrand for approaching a predator
integrand_ap_pr <- function(t, d, h) {
  sum_k <- 0
  for (k in -k_limit:k_limit) {
    term <- ((h - x + 2 * k * (h - d)) / (sqrt(2 * pi) * t^(3/2))) * exp(-(h - x + 2 * k * (h - d))^2 / (2 * t))
    sum_k <- sum_k + term
  }
  result <- exp(-mu * (h - x) - ((mu^2) * t / 2)) * sum_k * (0 + 1)
  return(result)
}
# Integrand for avoiding a predator
integrand_av_pr <- function(t, d, h, below_b_t) {
  sum_k <- 0
  for (k in -k_limit:k_limit) {
    term <- ((x - d + 2 * k * (h - d)) / (sqrt(2 * pi) * t^(3/2))) * exp(-(x - d + 2 * k * (h - d))^2 / (2 * t))
    sum_k <- sum_k + term
  }
  result <- exp(-mu * (d - x) - ((mu^2) * t / 2)) * sum_k * ((below_b_t * exp(-alpha * t)) + 1)
  return(result)
```

```
}

# Integrand for approaching a prey

integrand_ap_py <- function(t, d, h, below_b_t, above_b_t) {

  sum_k <- 0

  for (k in -k_limit:k_limit) {

    term <- ((h - x + 2 * k * (h - d)) / (sqrt(2 * pi) * t^(3/2))) * exp(-(h - x + 2 * k * (h - d))^2 / (2 * t))

    sum_k <- sum_k + term

  }

  result <- exp(mu * (h - x) - ((mu^2) * t / 2)) * sum_k * ((above_b_t * exp(-lambda * t)) +
((below_b_t * (1 - exp(-lambda * t)))) + 1)

  return(result)

}

# Integrand for avoiding a prey

integrand_av_py <- function(t, d, h, below_b_t) {

  sum_k <- 0

  for (k in -k_limit:k_limit) {

    term <- ((x - d + 2 * k * (h - d)) / (sqrt(2 * pi) * t^(3/2))) * exp(-(x - d + 2 * k * (h - d))^2 / (2 * t))

    sum_k <- sum_k + term

  }

  result <- exp(mu * (d - x) - ((mu^2) * t / 2)) * sum_k * (below_b_t + 1)

  return(result)

}


# Calculate the expected reward given thresholds d and h

expected_reward <- function(d, h, p, below_b_t, above_b_t) {

  E_ap_pr <- pracma::integral(integrand_ap_pr, xmin = 0, xmax = time_limit,

                             d = d, h = h, method = "Kronrod",

                             random = F, no_intervals = time_limit/time_step)

  E_av_pr <- pracma::integral(integrand_av_pr, xmin = 0, xmax = time_limit,
```

```r
                    d = d, h = h, below_b_t = below_b_t, method = "Kronrod",
                    random = F, no_intervals = time_limit/time_step)
  E_ap_py <- pracma::integral(integrand_ap_py, xmin = 0, xmax = time_limit,
                    d = d, h = h, below_b_t = below_b_t, above_b_t = above_b_t, method = 
"Kronrod",
                    random = F, no_intervals = time_limit/time_step)
  E_av_py <- pracma::integral(integrand_av_py, xmin = 0, xmax = time_limit,
                    d = d, h = h, below_b_t = below_b_t, method = "Kronrod",
                    random = F, no_intervals = time_limit/time_step)

  E_reward <- p * E_ap_pr + p * E_av_pr + (1 - p) * E_ap_py + (1 - p) * E_av_py
  return(E_reward)
}

# Optimize the thresholds d and h to maximize the expected reward
optimize_thresholds <- function(p, below_b_t, above_b_t) {
  d_range <- seq(-0.2, -2, by = -0.2)
  h_range <- seq(0.2, 2, by = 0.2)
  max_reward <- -Inf
  optimal_d <- NA
  optimal_h <- NA

  for (d in d_range) {
    for (h in h_range) {
      reward <- expected_reward(d, h, p, below_b_t, above_b_t)
      if (reward > max_reward) {
        max_reward <- reward
        optimal_d <- d
        optimal_h <- h
```

```
      }
    }
  }

  return(list(optimal_h = optimal_h, optimal_d = optimal_d, max_reward = max_reward))
}

# Function to run the optimization and store results of 200 iterations
run_optimization <- function(p) {
  values <- data.frame(reserves, b_t)
  thresholds <- data.frame(reserves, th_d, th_h)

  for (i in 1:200) {
    new_b_t_col_name <- paste0("b_t-", i)
    new_th_d_col_name <- paste0("th_d-", i)
    new_th_h_col_name <- paste0("th_h-", i)

    values[[new_b_t_col_name]] <- NA
    thresholds[[new_th_d_col_name]] <- NA
    thresholds[[new_th_h_col_name]] <- NA

    prev_b_t_col_name <- if (i == 1) "b_t" else paste0("b_t-", i - 1)

    for (j in 1:nrow(values)) {
      if (values$reserves[j] == 0) {
        values[[new_b_t_col_name]][j] <- 0
        thresholds[[new_th_d_col_name]][j] <- NA
        thresholds[[new_th_h_col_name]][j] <- NA
      } else {
```

```
        above_b_t <- if (j == 1) values[[prev_b_t_col_name]][j] else values[[prev_b_t_col_name]][j - 1]
        below_b_t <- if (j == nrow(values)) values[[prev_b_t_col_name]][j] else
values[[prev_b_t_col_name]][j + 1]

        list_thresholds_reward <- optimize_thresholds(p, below_b_t, above_b_t)
        thresholds[[new_th_d_col_name]][j] <- list_thresholds_reward$optimal_d
        thresholds[[new_th_h_col_name]][j] <- list_thresholds_reward$optimal_h
        values[[new_b_t_col_name]][j] <- list_thresholds_reward$max_reward
      }
    }
  }

  return(list(values = values, thresholds = thresholds))
}

# Run the optimization for different values of p
p_values <- seq(0.05, 0.95, 0.05)
results_list_2 <- list()

for (p in p_values) {
  results <- run_optimization(p)
  results_list_2[[paste0("p_", p, "_values")]] <- results$values
  results_list_2[[paste0("p_", p, "_thresholds")]] <- results$thresholds
}

# Extract the thresholds for reserves = 2 of the 200th iteration
plot_data_2 <- data.frame(p = p_values, th_d = NA, th_h = NA)

for (i in 1:length(p_values)) {
```

```
  p <- p_values[i]
  threshold_df <- results_list_2[[paste0("p_", p, "_thresholds")]]
  plot_data_2$th_d[i] <- threshold_df[3, "th_d-200"]
  plot_data_2$th_h[i] <- threshold_df[3, "th_h-200"]
}

# Plot Optimal thresholds vs Probability of predator
p2 <- ggplot(plot_data_2, aes(x = p)) +
  geom_line(aes(y = th_h, color = "Approach")) +
  geom_line(aes(y = th_d, color = "Avoid")) +
  labs(x = "probability of predator, *p*", y = "optimal thresholds") +
  scale_x_continuous(limits = c(0, 1), breaks = seq(0, 1, 0.1), expand = c(0, 0)) +
  scale_y_continuous(limits = c(-2, 2), breaks = seq(-2, 2, 0.5), expand = c(0, 0)) +
  scale_color_manual(values = c("Avoid" = "blue", "Approach" = "red")) +
  annotate("text", x = 0.4, y = 1.4, label = "Approach", color = "red") +
  annotate("text", x = 0.4, y = -1.2, label = "Avoid", color = "blue") +
  theme_classic() +
  theme(axis.title.x = ggtext::element_markdown(), legend.position = "none")


# Extract the expected values for reserves = 2 of the 200th iteration
plot_data_6 <- data.frame(p = p_values, exp_sur_time = NA)

for (i in 1:length(p_values)) {
  p <- p_values[i]
  exp_sur_time_df <- results_list_2[[paste0("p_", p, "_values")]]
  plot_data_6$exp_sur_time[i] <- exp_sur_time_df[3, "b_t-200"]
}
```

```
# Plot Expected survival time vs Probability of predator
p6 <- ggplot(plot_data_6, aes(x = p, y = exp_sur_time)) +
  geom_line(color = "black", linewidth = 1) +
  geom_point(shape = 2, size = 3, color = "black") +
  scale_x_continuous(name = "Probability of predator, *p*",
                     limits = c(0, 1),
                     breaks = seq(0, 1, 0.1),
                     expand = c(0, 0)) +
  scale_y_continuous(name = "Expected survival time",
                     limits = c(0, 65),
                     breaks = seq(0, 65, by = 5),
                     expand = c(0, 0)) +
  theme_classic() +
  theme(axis.title.x = ggtext::element_markdown(size = 12),
        axis.title.y = element_text(size = 12),
        axis.line.x = element_line(color = "black"),
        axis.line.y = element_line(color = "black"),
        panel.grid = element_blank(),
        axis.ticks = element_line(color = "black"),
        axis.ticks.length = unit(0.2, "cm"),
        legend.position = "none")
```

**Appendix C: Using a higher number of reserves levels**

The results presented in the main text remain consistent when increasing the number of possible reserve levels. Here, we used the same S3DM to maximize reproductive success, but with the maximum reserve level set to 10 and the dynamic programming iterative process run for 50 discrete time units. As shown in Figure C1, the same type of decreasing thresholds evident in the main text are observed, indicating a faster but less accurate approach response upon detecting another animal as the risk of predation increases.

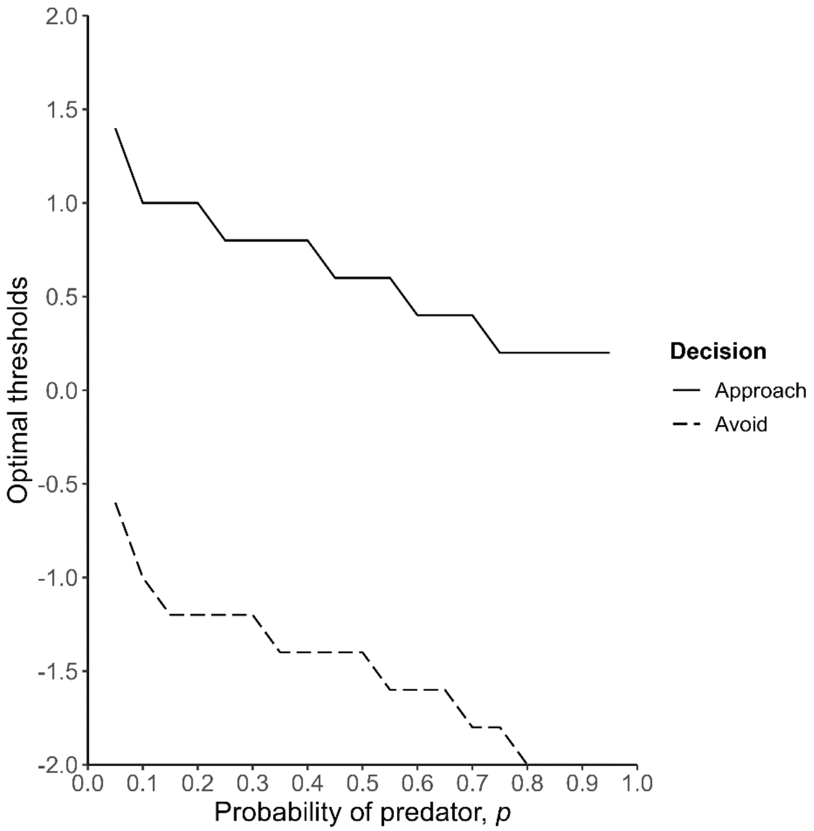


**Figure C1.** Optimal response thresholds for approach (solid line) and avoidance (dashed line) as a function of *p* in the S3DM when maximizing reproductive success. We use thresholds at a reserve level of 5. (Parameters: $\alpha$ = 0.2, $\lambda$ = 0.2, *c* = 6, *L* = 10.)

This is the code used to calculate the values for Figure C1:

```
library(pracma)

library(ggplot2)

library(dplyr)

library(ggtext)


# Define constants

alpha <- 0.2 # Speed of predator

lambda <- 0.2 # Speed of prey

mu <- 1 # Mean drift of information with time

x <- 0 # Starting point

time_step <- 0.1 # Time step for numerical integration
```

```
time_limit <- 10 # Time limit for numerical integration
k_limit <- 10  # Approximate the sum over the infinite k sum by running between limits of [-10, 10]



# Create initial reserves and t columns
reserves <- c(10:0)
b_t <- c(10:0)
th_d <- rep(NA, 11)
th_h <- rep(NA, 11)
L <- 10
c <- 6

# Create initial dataframe
values <- data.frame(reserves, b_t)
thresholds <- data.frame(reserves, th_d, th_h)

# Integrands

# Integrand for approaching a predator
integrand_ap_pr <- function(t, d, h) {
  sum_k <- 0
  for (k in -k_limit:k_limit) {
    term <- ((h - x + 2 * k * (h - d)) / (sqrt(2 * pi) * t^(3/2))) * exp(-(h - x + 2 * k * (h - d))^2 / (2 * t))
    sum_k <- sum_k + term
  }
  result <- exp(-mu * (h - x) - ((mu^2) * t / 2)) * sum_k * 0
  return(result)
}
# Integrand for avoiding a predator
```

```
integrand_av_pr <- function(t, d, h, below_b_t) {
  sum_k <- 0
  for (k in -k_limit:k_limit) {
    term <- ((x - d + 2 * k * (h - d)) / (sqrt(2 * pi) * t^(3/2))) * exp(-(x - d + 2 * k * (h - d))^2 / (2 * t))
    sum_k <- sum_k + term
  }
  result <- exp(-mu * (d - x) - ((mu^2) * t / 2)) * sum_k * ((below_b_t * exp(-alpha * t)) + (0 * (1 -
exp(-alpha * t))))
  return(result)
}
# Integrand for approaching a prey
integrand_ap_py <- function(t, d, h, below_b_t, above_b_t) {
  sum_k <- 0
  for (k in -k_limit:k_limit) {
    term <- ((h - x + 2 * k * (h - d)) / (sqrt(2 * pi) * t^(3/2))) * exp(-(h - x + 2 * k * (h - d))^2 / (2 * t))
    sum_k <- sum_k + term
  }
  result <- exp(mu * (h - x) - ((mu^2) * t / 2)) * sum_k * ((above_b_t * exp(-lambda * t)) +
(below_b_t * (1 - exp(-lambda * t))))
  return(result)
}
# Integrand for avoiding a prey
integrand_av_py <- function(t, d, h, below_b_t) {
  sum_k <- 0
  for (k in -k_limit:k_limit) {
    term <- ((x - d + 2 * k * (h - d)) / (sqrt(2 * pi) * t^(3/2))) * exp(-(x - d + 2 * k * (h - d))^2 / (2 * t))
    sum_k <- sum_k + term
  }
  result <- exp(mu * (d - x) - ((mu^2) * t / 2)) * sum_k * below_b_t
  return(result)
```

```
}

# Calculate the expected reward given thresholds d and h
expected_reward <- function(d, h, p, below_b_t, above_b_t) {
  E_ap_pr <- pracma::integral(integrand_ap_pr, xmin = 0, xmax = time_limit,
                         d = d, h = h, method = "Kronrod",
                         random = F, no_intervals = time_limit/time_step)
  E_av_pr <- pracma::integral(integrand_av_pr, xmin = 0, xmax = time_limit,
                         d = d, h = h, below_b_t = below_b_t, method = "Kronrod",
                         random = F, no_intervals = time_limit/time_step)
  E_ap_py <- pracma::integral(integrand_ap_py, xmin = 0, xmax = time_limit,
                         d = d, h = h, below_b_t = below_b_t,
                         above_b_t = above_b_t, method = "Kronrod",
                         random = F, no_intervals = time_limit/time_step)
  E_av_py <- pracma::integral(integrand_av_py, xmin = 0, xmax = time_limit,
                         d = d, h = h, below_b_t = below_b_t, method = "Kronrod",
                         random = F, no_intervals = time_limit/time_step)

  E_reward <- p * E_ap_pr + p * E_av_pr + (1 - p) * E_ap_py + (1 - p) * E_av_py
  return(E_reward)
}

# Optimize the thresholds d and h to maximize the expected reward
optimize_thresholds <- function(p, below_b_t, above_b_t) {
  d_range <- seq(-0.2, -2, by = -0.2)
  h_range <- seq(0.2, 2, by = 0.2)
  max_reward <- -Inf
  optimal_d <- NA
  optimal_h <- NA
```

```
  for (d in d_range) {
    for (h in h_range) {
      reward <- expected_reward(d, h, p, below_b_t, above_b_t)
      if (reward > max_reward) {
        max_reward <- reward
        optimal_d <- d
        optimal_h <- h
      }
    }
  }

  return(list(optimal_h = optimal_h, optimal_d = optimal_d, max_reward = max_reward))
}

# Function to run the optimization and store results of 50 iterations
run_optimization <- function(p) {
  values <- data.frame(reserves, b_t)
  thresholds <- data.frame(reserves, th_d, th_h)

  for (i in 1:50) {
    new_b_t_col_name <- paste0("b_t-", i)
    new_th_d_col_name <- paste0("th_d-", i)
    new_th_h_col_name <- paste0("th_h-", i)

    values[[new_b_t_col_name]] <- NA
    thresholds[[new_th_d_col_name]] <- NA
    thresholds[[new_th_h_col_name]] <- NA
```

```
    prev_b_t_col_name <- if (i == 1) "b_t" else paste0("b_t-", i - 1)


    for (j in 1:nrow(values)) {
      if (values$reserves[j] == 0) {
        values[[new_b_t_col_name]][j] <- 0
        thresholds[[new_th_d_col_name]][j] <- NA
        thresholds[[new_th_h_col_name]][j] <- NA
      } else if (values$reserves[j] == L){
        values[[new_b_t_col_name]][j] <- (L - c) + 1
      } else {
        above_b_t <- if (j == 1) values[[prev_b_t_col_name]][j] else values[[prev_b_t_col_name]][j - 1]
        below_b_t <- if (j == nrow(values)) values[[prev_b_t_col_name]][j] else
values[[prev_b_t_col_name]][j + 1]


        list_thresholds_reward <- optimize_thresholds(p, below_b_t, above_b_t)
        thresholds[[new_th_d_col_name]][j] <- list_thresholds_reward$optimal_d
        thresholds[[new_th_h_col_name]][j] <- list_thresholds_reward$optimal_h
        values[[new_b_t_col_name]][j] <- list_thresholds_reward$max_reward
      }
    }
  }


  return(list(values = values, thresholds = thresholds))
}


# Run the optimization for different values of p
p_values <- seq(0.05, 0.95, 0.05)
results_list_5 <- list()
```

```
for (p in p_values) {
  results <- run_optimization(p)
  results_list_5[[paste0("p_", p, "_values")]] <- results$values
  results_list_5[[paste0("p_", p, "_thresholds")]] <- results$thresholds
}

# Extract the thresholds for reserves = 5 of the 50th iteration
plot_data_5 <- data.frame(p = p_values, th_d = NA, th_h = NA)

for (i in 1:length(p_values)) {
  p <- p_values[i]
  threshold_df <- results_list_5[[paste0("p_", p, "_thresholds")]]
  plot_data_5$th_d[i] <- threshold_df[6, "th_d-50"]
  plot_data_5$th_h[i] <- threshold_df[6, "th_h-50"]
}


# Plot Optimal thresholds vs Probability of predator
p1 <- ggplot(plot_data_5, aes(x = p)) +
  geom_line(aes(y = th_h, linetype = "Approach")) +
  geom_line(aes(y = th_d, linetype = "Avoid")) +
  labs(x = "Probability of predator, *p*", y = "Optimal thresholds") +
  scale_x_continuous(limits = c(0, 1), breaks = seq(0, 1, 0.1), expand = c(0, 0)) +
  scale_y_continuous(limits = c(-2, 2), breaks = seq(-2, 2, 0.5), expand = c(0, 0)) +
  scale_linetype_manual(name = "Decision",
                     values = c("Approach" = "solid", "Avoid" = "longdash")) +
  theme_classic() +
  theme(axis.title.x = ggtext::element_markdown(size = 14),
      axis.title.y = element_text(size = 14),
```

```
axis.text = element_text(size = 12),

legend.title = element_text(size = 12, face = "bold"),

legend.text = element_text(size = 11),

legend.position = "right")
```
